# Cortico-basal-ganglia communication: Temporally structured activity for selective motor control

**DOI:** 10.1101/413286

**Authors:** Petra Fischer, Witold Lipski, Wolf-Julian Neumann, Robert Sterling Turner, Pascal Fries, Peter Brown, Robert Mark Richardson

**Affiliations:** Medical Research Council Brain Network Dynamics Unit, University of Oxford, OX1 3TH, Oxford, United Kingdom; Nuffield Department of Clinical Neurosciences, John Radcliffe Hospital, University of Oxford, OX3 9DU, Oxford, United Kingdom; Brain Modulation Laboratory, Department of Neurological Surgery, University of Pittsburgh School of Medicine, Pittsburgh, Pennsylvania, United States; University of Pittsburgh Brain Institute, Pittsburgh, Pennsylvania, United States; Department of Neurology, Campus Mitte, Charite - Universitaetsmedizin Berlin, Berlin, Germany; Department of Neurobiology, University of Pittsburgh, Pittsburgh, Pennsylvania, United States; Center for Neuroscience, University of Pittsburgh, Pittsburgh, Pennsylvania, United States; Center for the Neural Basis of Cognition, University of Pittsburgh, Pittsburgh, Pennsylvania, United States; Ernst Strüngmann Institute (ESI) for Neuroscience in Cooperation with Max Planck Society, 60528 Frankfurt, Germany; Donders Institute for Brain, Cognition and Behaviour, 6525 EN Nijmegen, Netherlands

**Keywords:** Subthalamic nucleus, gamma oscillations, spike coupling, communication through coherence, ECoG recording, gripping task

## Abstract

Despite the hard-wired structural connectivity of neural pathways, neural circuits allow context-dependent reactions to sensory cues by triggering the desired movement. Cortico-basal-ganglia circuits seem particularly important for flexible motor control as this is impaired in Parkinson’s disease (PD). We analysed subthalamic nucleus (STN) spike and cortical ECoG activity from PD patients performing a visually-cued hand grip task. Fast reaction times were preceded by enhanced STN spike-to-cortical gamma phase coupling irrespective of firing rate changes, suggesting a role of gamma coupling in motor preparation. STN spike timing was offset by half a cycle when comparing ipsilateral with contralateral movements. Additionally, cortical high-frequency activity increased more steeply within each gamma cycle at the sites that showed the strongest coupling with STN spikes. Cortico-basal-ganglia gamma coupling may thus help shape neural activity to facilitate selective motor control. The observation that this effect occurs independent of changes in mean firing rate has far-reaching implications.

**Highlights:**

- Fast RTs were preceded by enhanced STN spike-to-cortical gamma phase coupling
- STN spike probability was significantly modulated relative to the gamma cycle
- During ipsilateral movement, spikes were more likely at the opposite part of the cycle
- STN output may thus help shape cortical gamma for selective motor control

## Introduction

Interacting appropriately with the world is a strikingly complex feat if one considers the sheer infinite variety of movements that can be performed at any time. The basal ganglia seem to play a crucial role in motor control and learning (Kim et al., 2015; Piron et al., 2016; Yttri and Dudman, 2016) although their exact contribution to initiating or regulating movements is still unclear (Turner and Desmurget, 2010; Dudman and Krakauer, 2016). Recent work has shown that motor preparation involves a dynamic interplay between premotor cortical areas and portions of the ventral thalamus (Guo et al., 2017), which are tonically inhibited by basal ganglia output (Albin et al., 1989). The direct and indirect basal ganglia pathways are therefore well-placed to facilitate motor preparation by temporarily lowering inhibition but also by supporting suppression of unwanted movements. It is thus not surprising that both hypo- and hyperkinetic movement disorders such as Parkinson’s and Huntington’s disease can originate from basal ganglia degeneration (Albin et al., 1989).

Local field potential recordings from sites within the cortico-BG-thalamo-cortical (CBGTC) loop consistently show a relatively narrow band (~60-90 Hz) of gamma oscillations appearing at movement onset (Masimore et al., 2005; Brücke et al., 2008, 2012, 2013; Muthukumaraswamy, 2010; Jenkinson et al., 2013). Non-invasive MEG studies have also revealed a narrow-band gamma increase in primary motor cortex (M1), which quickly subsides during prolonged contractions (Muthukumaraswamy, 2010). This suggests that gamma synchrony may be important for movement initiation. Not only local power but also functional STN-to-M1 gamma connectivity increases during movement execution and is enhanced by dopaminergic medication (Litvak et al., 2012), which substantially improves a patient’s ability to move. LFP activity in this narrow band in the BG, specifically the STN, phase-precedes that in the motor cortex, suggesting that the basal ganglia modulate cortical activity (Williams et al., 2002; Litvak et al., 2012; Sharott et al., 2018). Previous studies of STN-to-M1 coupling have only examined STN activity at the population level using LFP signals, which are thought to reflect the changes in membrane potential shared across a large population of nearby neurons. These fluctuations are thought to arise predominantly from synaptic inputs (Buzsáki et al., 2012). Single-unit spiking activity, in contrast, reflects local output.

During visual processing, neuronal spikes that carry stimulus information to hierarchically higher cortical areas appear to be routed via gamma phase-synchrony, which can be flexibly enhanced or suppressed depending on which stimulus is attended (Bosman et al., 2012). These observations have been summarized in the Communication-through-Coherence theory (Fries, 2015), which also guides research on other essential cognitive functions such as working memory (Bastos et al., 2018).

Gamma power in the STN increases not only during movement initiation but also during fast stopping (Fischer et al., 2017), indicating that gamma synchrony may represent a mechanism for flexible information routing not only when processing sensory input but also for processing related to motor control. This idea is supported by observations that gamma oscillations reflect membrane potential fluctuations that establish short windows of relative excitation and inhibition (Hasenstaub et al., 2005; Gyorgi and Wang, 2012) providing brief opportunities for coordinated neuronal information transfer in the form of spikes.

To date, the majority of reports on neural correlates of movement preparation and execution have mainly focussed on firing rate changes (Anderson and Horak, 1985; Mink and Thach, 1991; Turner and Anderson, 1997; Seo et al., 2012; Arimura et al., 2013; Thura and Cisek, 2017). The precise timing of spikes, however, may be equally important, considering the considerable convergence (Oorschot 1996) and multiple recurrent connections within the CBGTC circuit. From a vast set of possible movement combinations, only specific muscles need to be activated at any one point while other “output units” should remain silent. Given the convergence of multiple excitatory and inhibitory inputs onto a single neuron and the non-linear membrane properties of neurons (Wilson and Kawaguchi, 1996; Deister et al., 2009), both of which apply for many neurons of the basal ganglia, the precise timing of an excitatory synaptic input substantially influences whether or not that input will elicit a spike.

Given that changes in STN spike rhythmicity and locking to cortical oscillations can occur before movement in the absence of increases or decreases in average firing rates (Lipski et al., 2017b), we hypothesize that the precise timing of STN spikes carries information about movement. As ECoG signals tend to show a very broad movement-related gamma power increase between 50-200 Hz, past research has focussed on the coupling of spikes to the *amplitude* of ECoG in the 50-200 Hz range (Shimamoto et al., 2013; Lipski et al., 2017b). Here, we explored whether we can detect STN spike-to-cortical narrow-band gamma *phase coupling* and whether the strength of such coupling correlates with behaviour.

Cortical ECoG and STN spikes were recorded in Parkinson’s disease patients undergoing deep brain stimulation surgery. Patients performed a visually-cued gripping task in which they squeezed a handgrip with their left or right hand within 2s after the Go cue for at least 100ms. Our analysis included 28 spike recordings from 12 patients (12 single-unit and 16 multi-unit recordings), from which spike-coupling to lower frequencies have been described previously (Lipski et al., 2017b).

Here, we focussed on analysing STN spike-to-cortical gamma phase coupling with four aims in mind. First, we will answer if STN-spike to cortical gamma phase coupling is strongest over hand regions of the motor cortex in this task. Second, we will test if the level of coupling is linked to variability in reaction times or movement kinematics. Third, we will test if the timing of STN spikes relative to cortical gamma oscillations is distinct for movements of contra- and ipsilateral hands. Finally, we will determine whether cortical sites that differ in how strongly they are coupled with STN spikes show different profiles of local cortical high-frequency activity, consistent with the interpretation that the effects reflect an impact of subcortical dynamics on cortical function.

## Results

### Gripping performance

Average reaction times from the Go signal to the grip onset were 0.53 ± (SD) 0.21s and 0.55 ±0.18s for contra- and ipsilateral grips, respectively (contra-ipsi: P = 0.435, t_27_ = −0.8). The average grip duration was 0.98 ±0.47s and 0.98 ±0.43s (contra-ipsi: P = 0.904, t_27_ = −0.1). The mean variability (SD) of RTs within each patient was 0.19 ±0.09s and 0.23 ±0.12s (contra-ipsi: P = 0.050, t_27_=-2.1). This variability in RTs allowed us to compare the extent of spike-to-gamma phase coupling between trials with fast and slow RTs. To this end, the data were grouped into trials with RTs below the median and trials with RTs above the median. The average RTs of these two sets were 0.39 ±0.16s and 0.68 ±0.27s for grips contralateral to the recorded STN and 0.38 ±0.12s and 0.72 ±0.26s for ipsilateral grips. The delay between the cue that instructed patients whether to perform the grip with the right or left hand and the Go signal did not differ significantly between short and long RT trials (difference in delay between long vs. short RT trials = 4ms, P= 0.116).

We also performed median-splits according to two other movement parameters: grip peak force (contralateral grips: 91 ±65 N vs. 104 ±71 N; ipsilateral grips: 80 ±56 N vs. 93 ±61 N) and peak yank (contralateral grips: 0.39 ±0.30 N/ms vs. 0.50 ±0.36 N/ms; ipsilateral grips: 0.33 ±0.23 N/ms vs. 0.43 ±0.29 N/ms).

### Topography of STN spike-to-cortical gamma coupling

ECoG strips were distributed predominantly over precentral and parietal areas and were always located in the same hemisphere as the recorded STN (Figure 1A). Cortical power in the 12-30 Hz beta and the 60-80 Hz gamma frequency bands was modulated at movement onset (−-0.1 – 0.4s around grip onset) in bipolar contact pairs situated predominantly in primary motor and sensory areas (Figures 1C+D), which is in line with past ECoG studies (Ohara et al., 2000; Pfurtscheller et al., 2003). The choice of 60-80 Hz for the gamma frequency band was based on reports of significant gamma coupling in this range between STN LFP and cortical MEG (Figure 5 and 6 from Litvak et al., 2012). The bipolar contact pairs with the strongest cortical gamma phase coupling to STN spikes during contralateral gripping were concentrated in lateral precentral gyrus (Figure 1E), a site, which corresponds strikingly well to a hotspot of 60-90 Hz gamma synchronization detected in MEG studies (Figure 1B, adapted from Cheyne et al., 2008). These sites, as well as the set of sites that showed the highest coupling during ipsilateral gripping were selected for further analyses. The spatial focus of coupling during contralateral gripping becomes even clearer when only the sites are displayed for which movement-related STN-spike-to-cortical gamma phase coupling reached significance (Figure 1F). The sites that showed significant coupling during ipsilateral gripping were more widely dispersed. The significance tests in Figure 1F show that gamma coupling was higher relative to a pre-movement period and relative to shuffled data, confirming that the increase exclusively occurred at movement onset and was not already present at rest.

**Figure 1.**
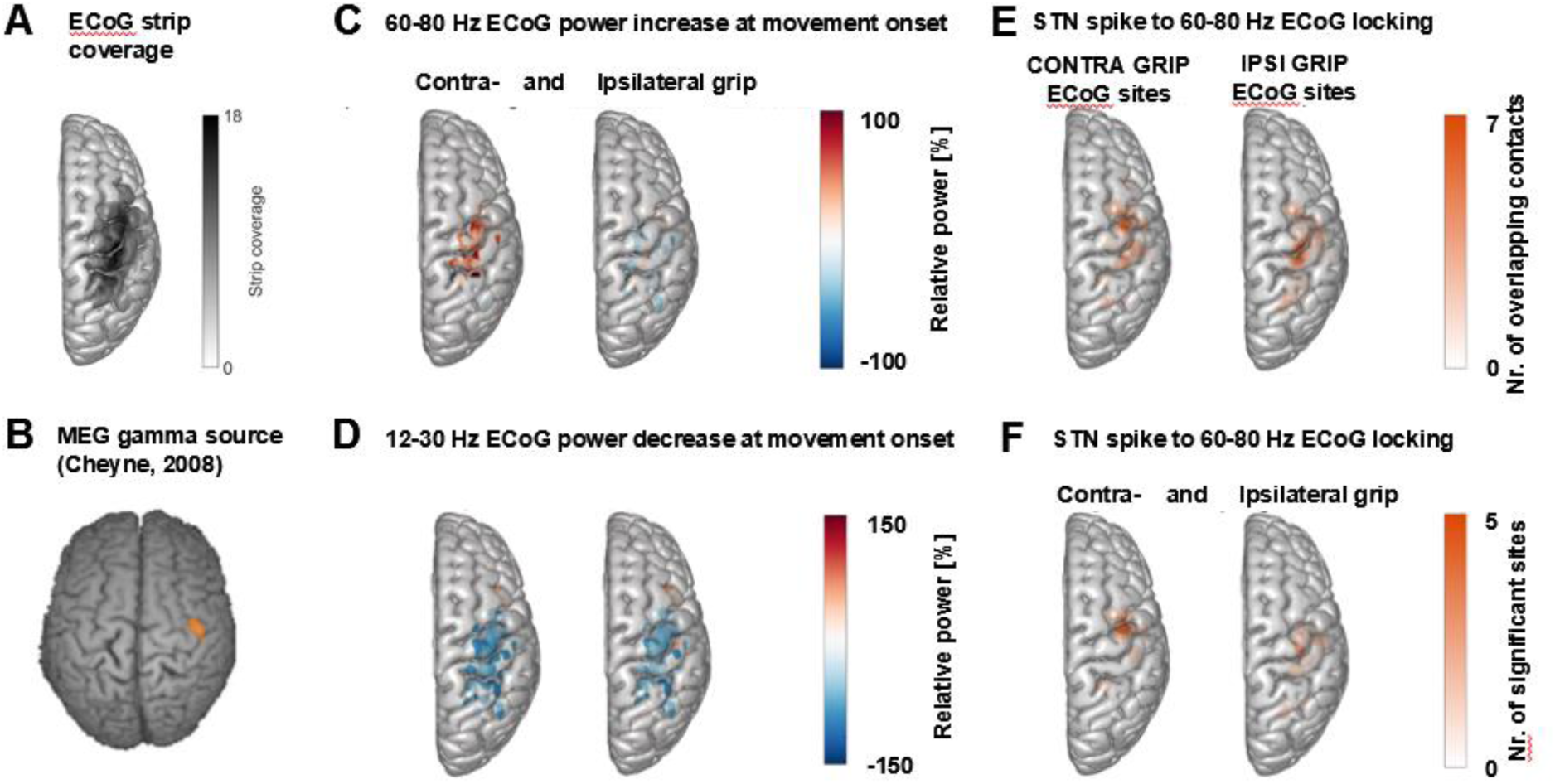
Cortical topography of STN spike-to-cortical gamma locking and power changes. **A** ECoG strips were distributed mainly over precentral and parietal areas. ECoG coordinates from recordings performed in the left hemisphere were flipped in the x-axis to allow averaging across recordings performed in the left and right hemisphere. The maximum number of overlapping recordings was 18 (dark areas). **B** Cheyne et al. (2008) showed that gamma oscillations at the onset of finger movements are focal to lateral motor cortex in MEG recordings (adapted with permission). **C** Gamma power increased most strongly over motor and somatosensory cortical areas during contralateral gripping. **D** Beta power decreased in spatially more widespread areas. **E** The left and right plots show the two sets of sites that showed the highest 60-80 Hz coupling during contralateral gripping and ipsilateral gripping, respectively. Each set includes one single site per recorded hemisphere (n=28). These sites were chosen for further analyses. **F** as **E** but reduced to the sites showing significant gamma coupling −0.1 – 0.4s around movement onset (α=0.1 to show the location of a larger number of contacts). If several channels of one recording were significant, only the channel with the highest PLV was included. The contacts that showed significant gamma PLV during contralateral gripping were concentrated over precentral cortex. This hotspot corresponds well with the 60-90 Hz gamma source localized in MEG studies (Cheyne et al., 2008) shown in **B**). The significant contacts for ipsilateral gripping were scattered over a wider area.

**Figure 5.**
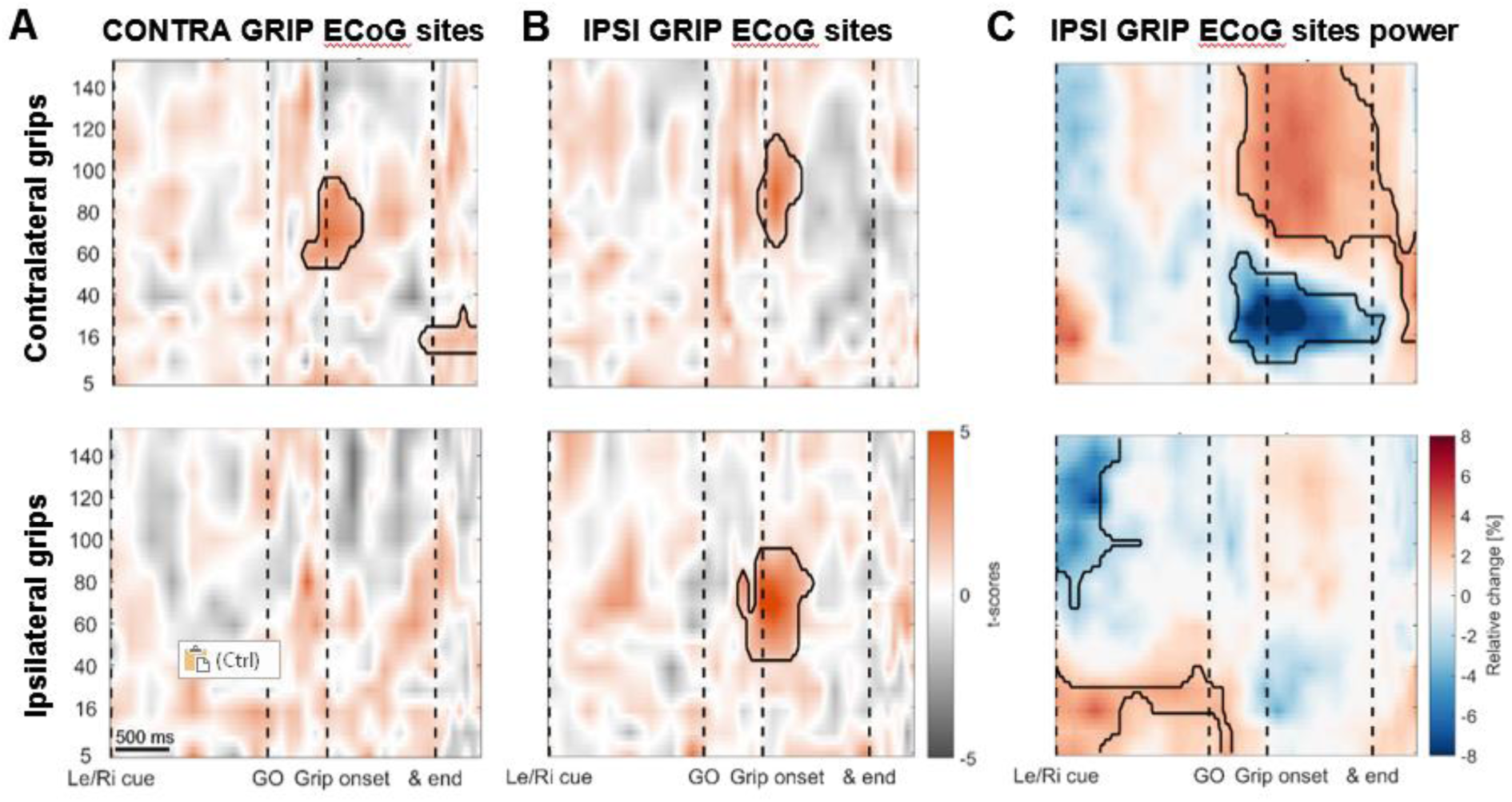
During ipsilateral gripping, STN spikes can be locked to gamma recorded from different cortical sites. Coupling strength is presented for two different sets of ECoG sites: One set of contacts with highest gamma PLV during contra-**(A)** and the other one with highest gamma PLV during ipsilateral gripping **(B)**. The leftmost column displays the same plots as shown in Fig. 2 to enable a direct comparison. **B** Significant locking between STN spikes and cortical sites with the highest 60-80 Hz PLV during ipsilateral gripping (bottom row). Note that in these sites, locking also occurs for contralateral grips (top row) although the gamma frequency is slightly higher as when calculated for the other set of ECoG sites shown in **A**. **C** In IPSI GRIP ECoG sites, gamma power increased again significantly only during contralateral gripping. Note that in response to the Left/Right cue, which announced an ipsilateral movement, beta power was significantly elevated up until the Go signal while >60 Hz power was relatively suppressed. Significant clusters (encircled in black) were obtained with a cluster-based permutation procedure to correct for multiple comparisons (p < 0.05).

**Figure 6.**
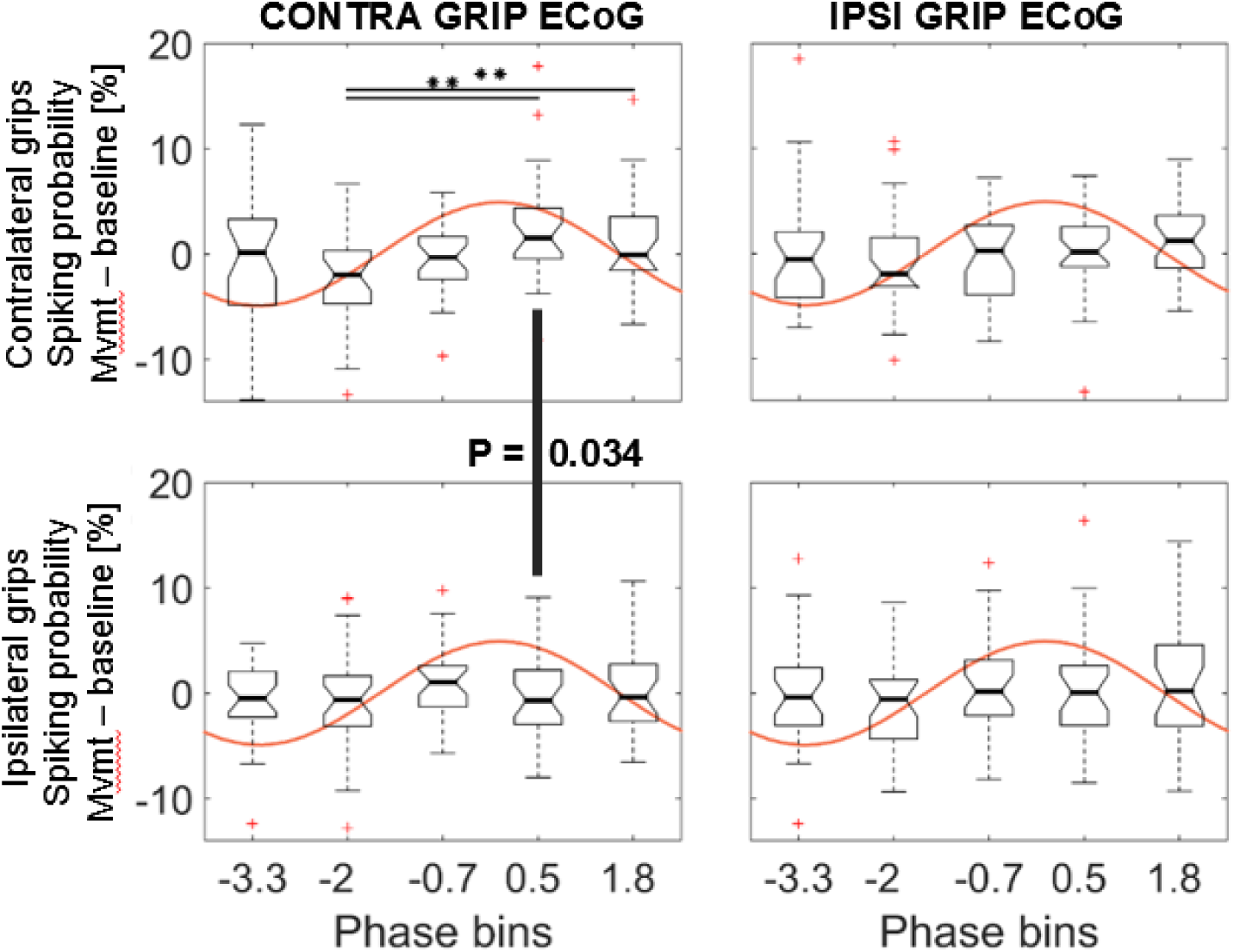
STN spiking probability is modulated relative to the phase of cortical gamma cycles. The five non-overlapping phase bins were set such that one bin (BIN4) was centred around the average preferred gamma phase during contralateral gripping across all recordings. The top left plot shows how spiking probabilities during contralateral grip onset co-modulate with the gamma phase.* show FDR-corrected significant differences between phase bins. Spiking was relatively suppressed in BIN2, which was centred around 2 rad before the gamma peak. The spiking probability in BIN4 was higher in contralateral grip trials than during ipsilateral gripping (P = 0.034, uncorrected for multiple comparisons). The bottom row shows ipsilateral grip trials and the right column shows the same analysis with the gamma phase extracted from ECoG sites showing the highest PLV during ipsilateral gripping. A significant interaction and main effect of bins was only found for the ANOVA based on gamma from ECoG sites that showed the highest PLV during contralateral gripping (left column). The central mark of the box plots displays the median and edges show the 25^th^and 75^th^ percentile. Whiskers show the 1.5*interquartile range and outliers (red crosses) are data points beyond this range.

### Movement-related STN spike-to-cortical gamma coupling

To evaluate changes in coupling across the whole task period and across multiple frequencies at the group-level, we employed a novel procedure that ensured that the time-frequency plots depict changes in the data locked to each task event. Each trial began with a Left/Right cue that was followed by a GO signal, movement onset and movement offset. Intervals between events were divided into equidistant points for each trial so that the variability of these intervals across trials and subjects did not affect the accuracy of the data alignment around important task events. The windows centred around these points were scaled such that each window contained the same numbers of spikes that was used to calculate the phase-locking value (PLV) for this point (see Methods). We pre-selected the set of ECoG contacts with the highest PLV between 60-80 Hz over a −0.1 – 0.4s period around movement onset (Figure 1E). Accordingly, STN firing was consistently phase-locked to cortical gamma at movement onset in these sites across all recordings (Figure 2A, n = 28). The significant cluster shows that coupling strength was higher than that obtained from a permutation distribution, which was generated by shuffling the association between STN spike trains and cortical LFP gamma phase time courses across trials, while keeping the respective individual trials intact. The cluster highlights that in the selected sites, above-chance coupling was relatively constrained to a narrow window of gamma band around movement onset instead of being widespread, which would have been possible in principle, despite the selection criterion. During ipsilateral gripping, no significant clusters in coupling between STN spikes and the phase of this pre-selected set of ECoG sites were found (Figure. 2A, bottom row). Figure S1 shows the spike-triggered average of the 60-80 Hz filtered ECoG signal of three example recordings demonstrating clear locking around contralateral grip onset but not in a pre-movement period.

**Figure 2.**
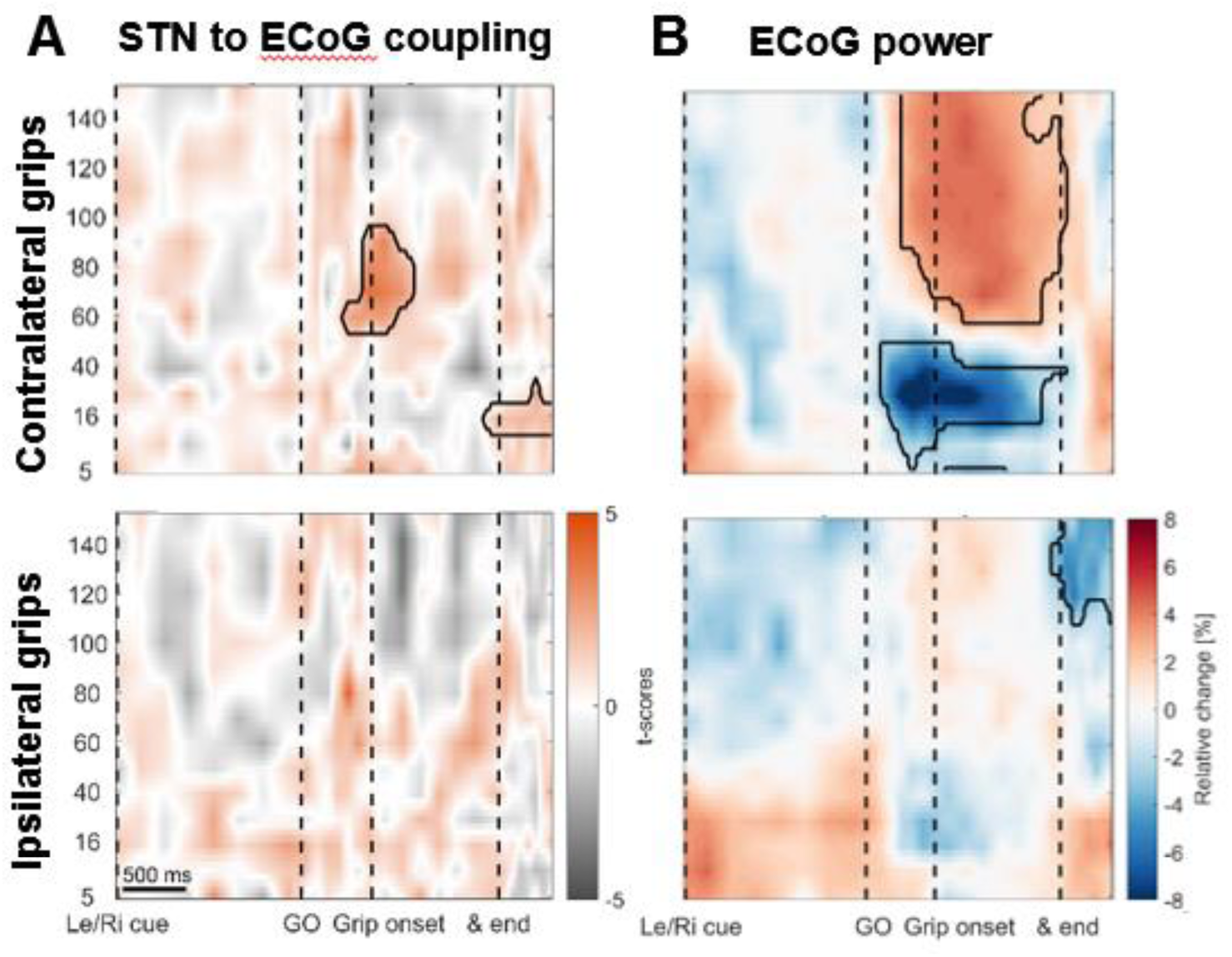
STN spike-to-ECoG phase coupling and ECoG power during visually cued gripping. The top and bottom row show contra- and ipsilateral grip trials, respectively. To get comparable estimates of the coupling strength around each of the key trial events, we implemented an event-locked variable-window width PLV estimation procedure: Each interval between neighbouring task events was subdivided into equidistant points, which served as centre for a window within which the PLV was computed. The windows varied in width to encompass the same number of spikes for all time points for one recording. **A** The black outlines show significant clusters obtained with a cluster-based permutation procedure for multiple comparison correction (n=28, p<0.05). Significant 60-80 Hz spike-to-cortical gamma phase coupling occured at contralateral grip onset. The same cells also locked to beta after movement offset when cortical beta power rebounded. During ipsilateral gripping (bottom row), no significant clusters were found. **B** During contralateral gripping, cortical gamma power increased while beta power was significantly suppressed. Ipsilateral gripping resulted in relatively reduced >120 Hz power after the grip was released.

As will be noted, the pre-selection of ECoG channels was supported by several independent findings. One is already apparent in Figure 2A; the same pre-selected contralateral ECoG sites were phase-locked to unit activity not only in the gamma band during movement but also to beta oscillations after movement completion when cortical beta power rebounded (Figure 2B). The left column in Figure S2 (Examples 1-3) shows examples of individual cells that coupled both to gamma at movement onset and beta after movement offset.

As already seen in the cortical surface plots, cortical gamma power increased significantly only during contralateral gripping, which was accompanied by a movement-related beta decrease (Figure 2B).

### Firing rates did not change on the group level

Next, we investigated event-related changes in firing rate and three other firing characteristics that could capture changes in spike patterns and have previously been reported in the context of spike-to-LFP coupling (Rule et al., 2017): the interspike interval coefficient of variation (ISI CV), percentage of bursting (defined as ISIs < 10ms) and the mode of the ISI (see Methods).

At the group level, no significant change in mean firing rate or proportion of bursting was found across the analysis interval (Figure 3). During contralateral gripping, the ISI CV was significantly reduced shortly after the Left/Right cue relative to a baseline obtained from the whole recording (Figure 3). Although Figure 3 suggests that a difference may have been present before the cue, Figure S3 shows that the drop occurred after the initial cue.

In ipsilateral grip trials, the ISI CV also was significantly reduced before and also during the movement. Additionally, the ISI mode decreased when ipsilateral grips were released, denoting shorter ISIs.

**Figure 3.**
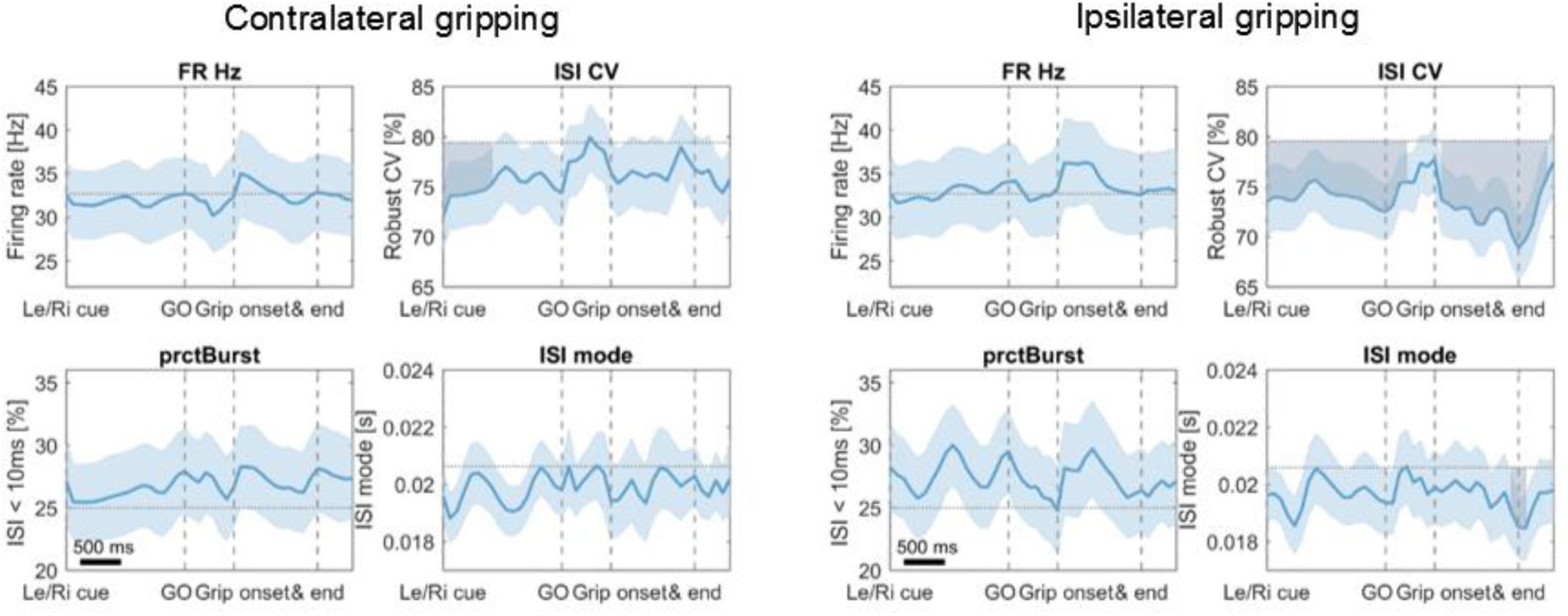
Firing characteristics during contra- and ipsilateral gripping. Values were compared against the baseline activity obtained from the whole recording (horizontal dotted line) using a cluster-based permutation procedure to correct for multiple comparisons similar as for the time-frequency plots. Significant deviations from baseline are shown as shaded areas in gray. In contralateral grip trials, the interspike interval CV was significantly reduced after the initial Left/Right cue onset. Although here it seems that the difference already was present when the cue was shown, Figure S3 shows that this effect was locked to cue onset. In ipsilateral grip trials, the ISI CV also was significantly reduced after the initial cue and in this case also during gripping. Additionally, a drop in ISI mode was observed at ipsilateral grip offset. Error bars show standard errors. Firing characteristics were computed similar as in Rule et al (2017). FR = firing rate in Hertz, ISI CV = Coefficient of variation of interspike intervals, prctBurst = percentage of bursting (defined as ISI < 10ms), ISI mode = mode of the interspike interval distribution.

As noted above, firing rates did not change significantly on the group level, but the 28 recordings were also assessed individually with cluster-based permutation tests as shown for the peri-stimulus time histograms in Figure S2. Cells were classified as increasing or decreasing their firing rates if a significant cluster occurred between the Go cue and the movement offset. Significant changes were tested against the baseline firing rate obtained from the whole recording.

21% of all recorded STN units significantly increased and 32% decreased their firing rates during contralateral movement while the rest (47%) showed no significant change. Three of the cells that increased their firing rates during gripping also showed a significant decrease either before or after the movement (Figure S2, Example 5). In 64% of all recordings, the firing behaviour did not differ between contralateral and ipsilateral. In the remaining recordings, firing behaviour differed between contralateral and ipsilateral gripping. Importantly, gamma coupling could be found in all types of cells irrespective of whether firing rates remained stable, increased or decreased (Figure S2). We evaluated this further by computing a correlation between firing rate changes at movement onset and coupling strength, which confirmed that no relationship was present (Pearson’s r = 0.01, P = 0.969, see Figure S4).

### Stronger spike-to-gamma coupling precedes faster reaction times

To test if differences in motor performance such as differences in reaction time, peak yank or peak force were linked to differences in gamma coupling, we median-split all trials and calculated coupling strengths separately for the two subsets. This analysis revealed that faster reaction times were preceded by higher STN spike-to-cortical LFP gamma coupling. The difference was present already at the time of the Go signal (Fig. 4A), which occurred on average 0.39s before grip onset in fast trials and 0.69s in trials with slow RTs.

**Figure 4.**
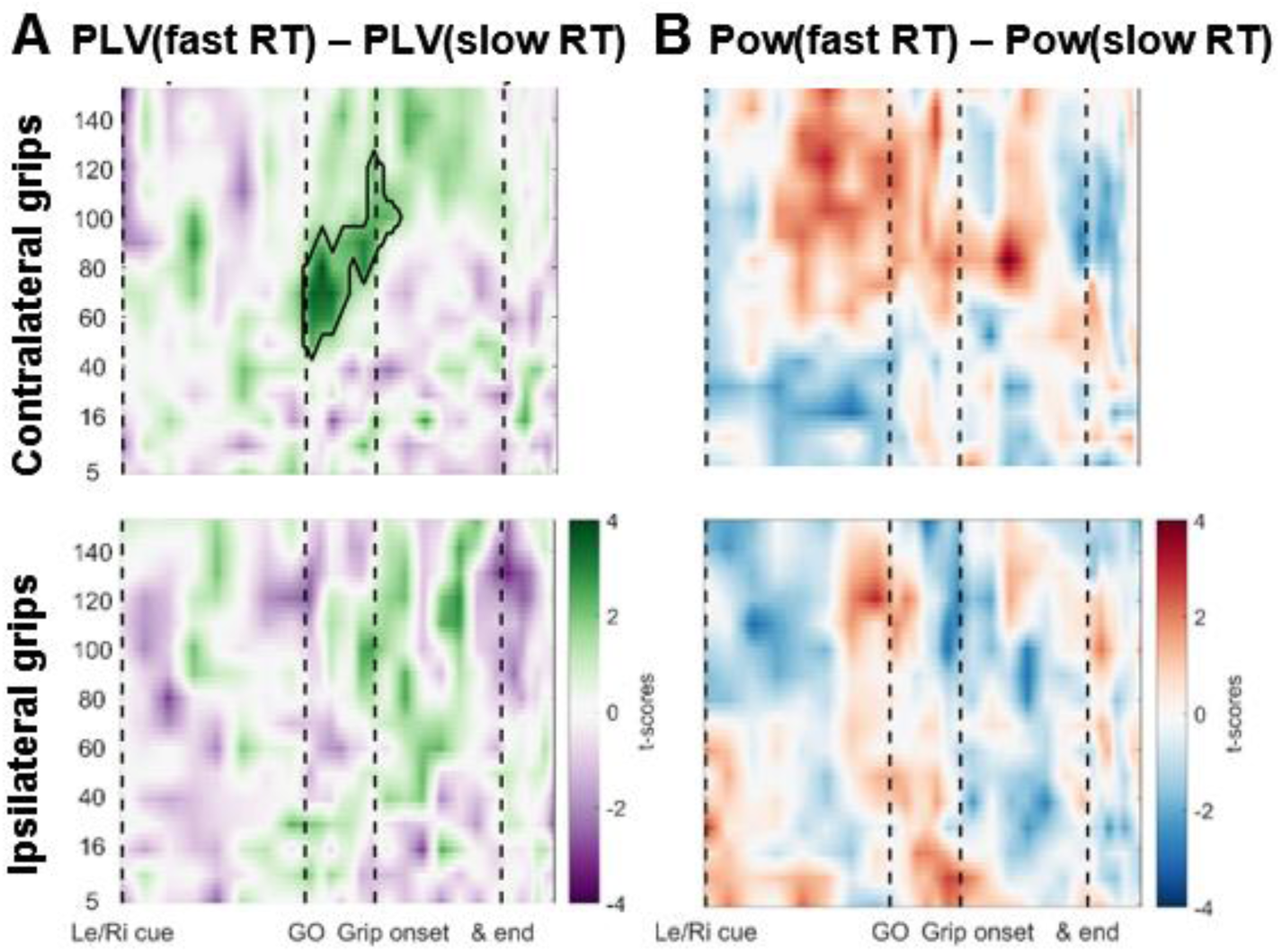
Coupling was significantly higher when reaction times were faster. **A** The green cluster shows that in those trials, in which RTs were fast as defined by a median split, gamma locking in the contralateral hemisphere was already significantly higher immediately after the Go signal, about 500ms before movement onset. **B** Average cortical gamma power tended to be higher and pre-Go signal beta power lower in contralateral grip trials with faster reaction times, however, these differences did not survive cluster-based multiple comparison correction.

The windows for calculating PLVs spanned on average −0.16:0.16s around each point. The precise window length at the Go signal was 306 ±68ms and 310 ±75ms for fast and slow RT trials. Thus, even in trials with short RTs, the movement onset fell outside the window used for calculating PLVs at the time of the Go signal, where a significant RT effect on coupling already was present (t_27_ = 3.2, P = 0.004). Hence, the difference in coupling cannot be explained merely by earlier movements in trials with short RTs. The difference disappeared at grip onset, which shows that coupling accompanying the movement itself was comparable irrespective of whether RTs leading up to the movement were fast or slow.

Local cortical gamma power appeared to be higher as well, although the effect was not significant (Fig. 4B). The effect of gamma coupling on RTs was specific to contralateral gripping. Firing characteristics between trials with fast and slow RTs again did not differ at the group level (Figure S5). When trials were split according to peak yank or peak grip force, no significant effects were found (Figure S6).

### Gamma coupling appears during ipsilateral gripping in different cortical sites

We next selected ECoG sites that showed the highest gamma coupling with STN spikes during ipsilateral grip trials and also found that this coupling exceeded that obtained by a permutation distribution (Figure 5B bottom). Note though, that these sites were spatially more widely distributed and tended to be located posterior to motor cortex (Figure 1E right, labelled IPSI GRIP ECoG sites). In these sites, gamma coupling was also present at contralateral grip onset (Figure 5B top), although the coupling frequency appears to be slightly higher than for the set of ECoG sites where coupling was largest for contralateral grips. Gamma power increased and beta power decreased significantly again only for contralateral grip trials (Figure 5C). Notably, if the initial cue indicated an ipsilateral grip, beta power was significantly elevated until the Go signal. During roughly the same time period, >60 Hz power was significantly reduced (Figure 5C).

### STN spiking probability is modulated by cortical gamma phase

Phase-locking values for individual pairs of STN spike and cortical ECoG recordings as a measure of coupling strength do not provide a measure of the consistency of the preferred phase across recordings. As only the length of the average phase vector is used to calculate PLVs, reflecting how strongly bundled or spread the phase values are, information about the preferred phase is lost. We next tested whether, across recordings, spikes were consistently more probable at certain points of the gamma cycle and less probable at others. Conversely, it would be unlikely to find a consistent phase preference and significant modulation of spike probabilities if high PLV coupling was a by-product of analysis choices (e.g., the pre-selection of ECoG contacts).

To examine if the spiking probability was consistently increased or decreased at certain phases of the gamma cycle, we subdivided the continuous cortical gamma signal into five non-overlapping phase bins (width = 0.4π) to calculate the percentage of spikes falling into these bins. Five bins were chosen to allow for a resolution that is high enough to detect differences but still results in a limited number of multiple comparisons. The binning was preceded by a phase-flipping procedure to standardize the phase polarity of the bipolar recordings (see Methods and Figure S7 A).

Spiking probabilities in each bin were computed within a −0.1 – 0.4s window around movement onset, and to ensure that the effects are movement-onset-specific, these probabilities were normalized by baseline spike probabilities, which were computed within a same sized window starting 3s before movement onset.

The centres of the five non-overlapping phase bins were set such that one bin was centred around the average preferred gamma phase during contralateral gripping across all recordings. The 5 (*bins*) x 2 (*effector side: contralateral vs. ipsilateral*) ANOVA resulted in a significant main effect of the factor *bins* (F_4,108_= 3.3, P = 0.022) and a significant interaction (F_4,108_= 2.5, P = 0.044) and no main effect of effector side (F_1,27_= 0.02, P = 0.877). The significant main effect of bins shows that those sites that were pre-selected for highest coupling irrespective of the preferred phase, have shared preferred and non-preferred phases. Figure 6 (left column) shows that spiking probabilities were significantly modulated during contralateral gripping: Spiking probabilities in BIN4+5 were significantly higher than in BIN2 (Wilcoxon signed-rank test, P = 0.001 and t_27_=-3.0, P = 0.005). Only the ANOVAs for gamma obtained from ECoG sites that showed the highest coupling during contralateral gripping were significant. Spike probabilities for gamma obtained from the other set of sites are shown in the right column of Figure 6.

The significant interaction suggests that the spiking probability in some bins differed between contralateral and ipsilateral grip trials. Pairwise comparisons showed that during contralateral grip trials, the spiking probability was enhanced in BIN4 (centred at 0.5 rad of the gamma cycle) although this did not survive multiple-comparison correction (P=0.034 uncorrected).

We also shifted the bin centres slightly to demonstrate that the above configuration is not the only one producing such results. The alternative binning configuration (the first bin started at 0.5π instead of 0.24π as in the analysis above) resulted in similar effects in the ANOVA (Figure S7 B, significant main effect bins: F_4,108_= 4.6, P = 0.004, main effect effector n.s.: F_4,108_= 0.03, P = 0.868, significant interaction: F_1,27_= 2.8, P = 0.035). The differences in spiking probabilities between contra- and ipsilateral grip trials were stronger in this configuration and survived multiple-comparison correction for BIN 5 (BIN5 at 0.9 rad: P = 0.003; BIN3 at −1.6 rad P = 0.040, n.s. after multiple-comparison correction). When looking at individual cells instead of the average, the largest three differences in spike probability between BIN5 and BIN3 were 21, 21 and 19% indicating that the probability for spikes to occur in BIN3 could be up to 21% lower than for spikes in BIN5.

This distinct modulation of spike probability according to the phase of gamma may provide a mechanism that enhances cortical gamma oscillations, which were more pronounced during contralateral gripping (Figures 2+5). In the next section, we will test if STN spikes were systematically offset relative to the cortical gamma phase during contra-versus ipsilateral gripping.

Because spike probabilities were only significantly modulated relative to gamma from ECoG sites that showed the highest coupling during contralateral gripping, although both sets of ECoG sites showed a gamma power increase, we wanted to investigate local cortical activity more closely. In the final section, we will thus test if the temporal profile of local excitability that fluctuates within each gamma cycle, differs between the two sets of sites.

### The preferred gamma phase of STN spikes differs between contra- and ipsilateral movements

To test if the timing of STN spikes relative to the cortical gamma phase systematically differed between contra- and ipsilateral grip trials, we extracted the gamma phase from ECoG sites that showed the highest coupling during contralateral gripping and computed the average, or “preferred”, phase coinciding with STN spikes for both trial types. To exclude recordings where estimation of the average phase estimate was unreliable, we started conservatively by using only recordings in which gamma coupling was significant at movement onset (shown in Figure 1F). We found in this subset (n=11) that the preferred gamma phase differed significantly between contra- and ipsilateral grip trials by ~210° (mean offset −2.5 rad, 95% CI = [−3.9, −1.1], Figure 7A). A more relaxed selection included all recordings in which the pairwise phase consistency value (a measure of unbiased coupling strength) exceeded zero (see Methods). In the larger set (n=23), which is potentially more representative although individual phase estimates may be more noisy, the preferred phase was offset by approximately 180° (Figure 7B, mean offset 3.1 rad, 95% CI = [2.1, 4.1]). P-values derived from a V-test assessing directionality towards an offset of 180° showed that the strength of the phase difference effect was strongest in a 500ms sliding window centred around 0.15 after movement onset, i.e. within the −0.1 – 0.4s window around movement onset (Figure 7C).

**Figure 7.**
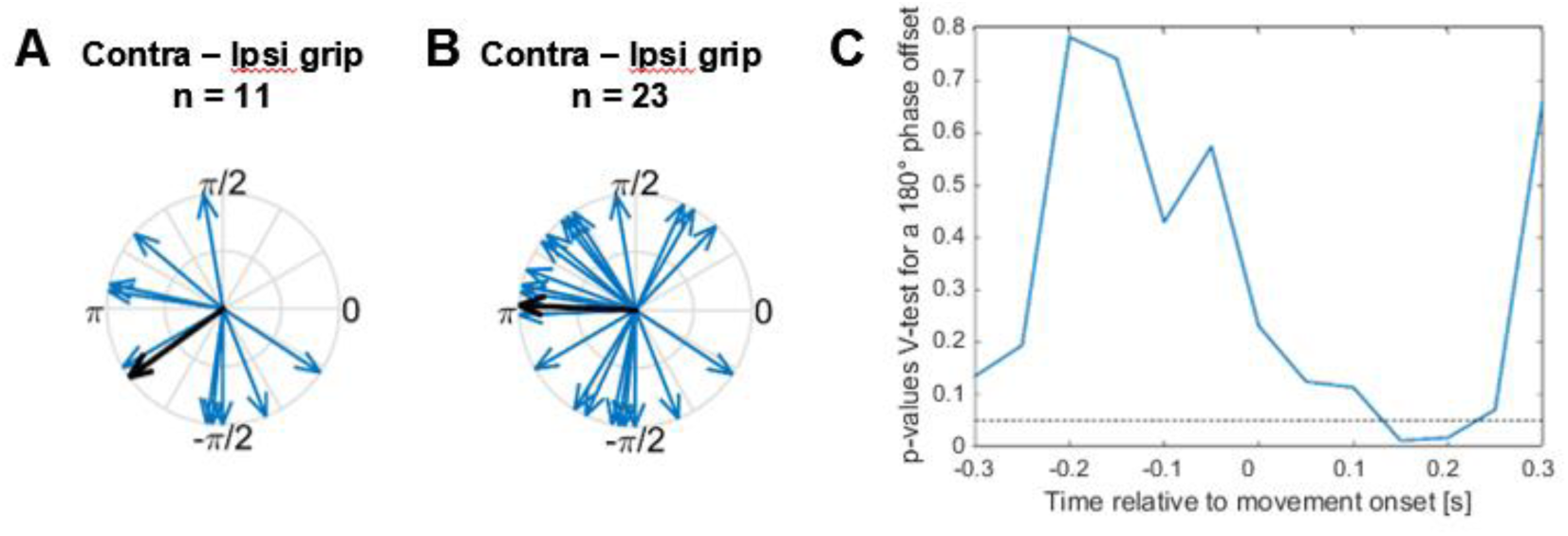
Systematic offset of STN spike timing relative to cortical gamma between contra- and ipsilateral gripping. **A** STN spikes of recordings with significant gamma coupling (n = 11) coincided with cortical gamma phases that were on average nearly opposite when comparing contra- and ipsilateral gripping (mean offset = −2.5 rad, 95% CI = [−3.9, −1.1]). **B** When including more recordings (all recordings where PPC > 0, n = 23), the offset was very close to 180° (mean offset =3.1 rad, 95% CI = [2.1, 4.1]). The gamma phase was extracted from ECoG sites that showed the highest coupling during contralateral gripping. **C** P-values derived from a V-test assessing directionality towards an offset of 180° (n = 23) show that the effect of the offset is strongest in a 500ms sliding window centered around 0.15 after movement onset, i.e. in a window −0.1 – 0.4s around movement onset.

### Cortical high-frequency activity increases faster in sites that show strongest gamma coupling during contralateral grips

In this final section, we will investigate whether the temporal profile of local excitability nested within the gamma cycle differed between cortical sites with high STN spike-to-cortical-gamma phase coupling and sites that were less coupled. To test this, we first computed the cortical high-frequency activity (HFA: rectified >300 Hz activity). Then we partitioned the 60-80 Hz gamma phase vector, which was extracted from the same signal, into non-overlapping phase bins for which the average HFA activity was computed. This resulted in an HFA time course that resembled the trough and peak of the gamma cycle (Figure 8A).

**Figure 8.**
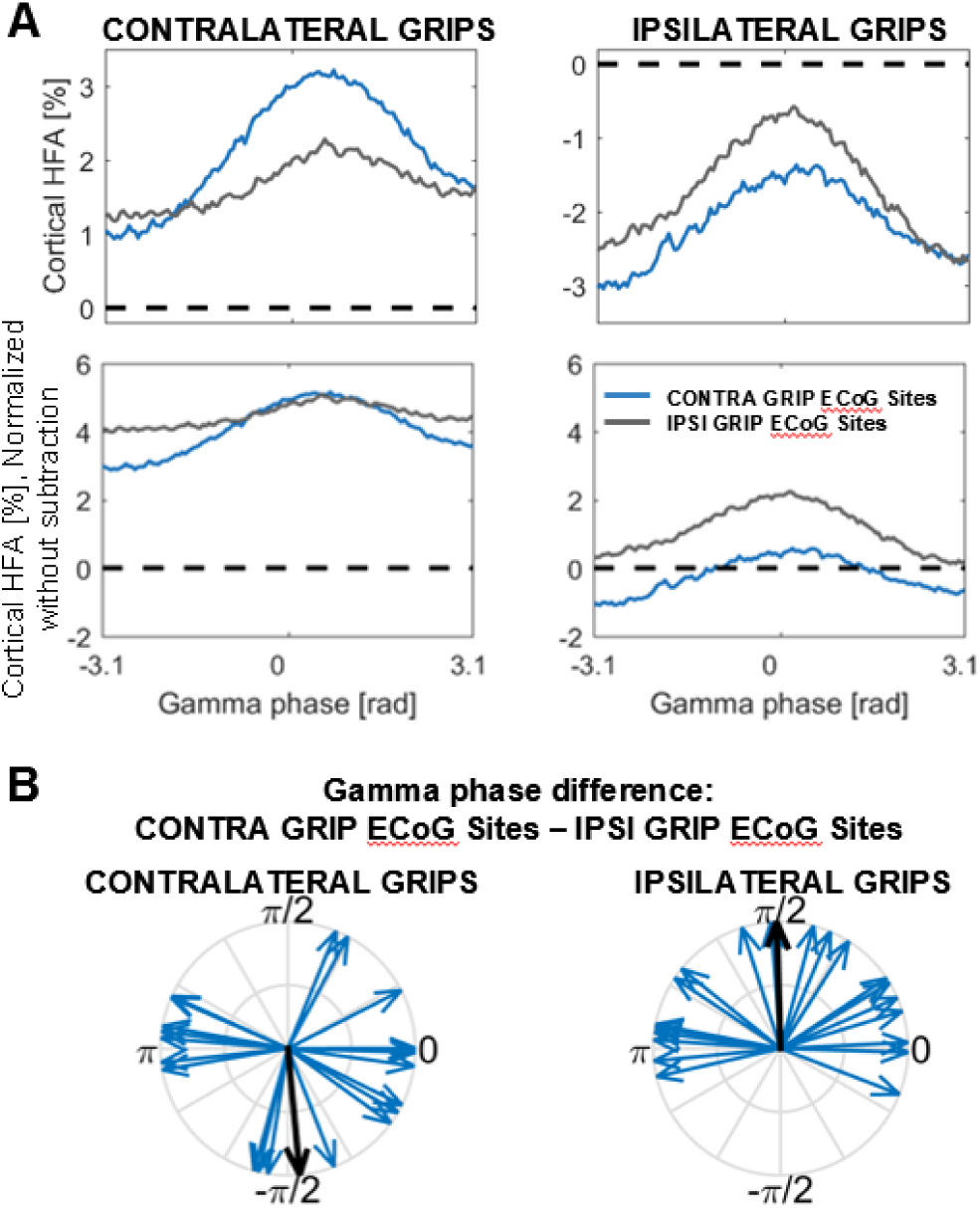
Cortical high-frequency activity (HFA) during contra- and ipsilateral gripping (left and right column respectively). **A** The blue and gray line show HFA from the two sets of ECoG sites with highest gamma PLV during contra- and ipsilateral gripping, respectively (n=20 as 8 recordings were excluded because the ECoG site was the same for both grip types). The data was normalized first by the mean of the recording. To remove any offset between the sets of sites, each line was subtracted by the mean of the two lines sharing the same colour (top row). The row below shows the normalized data without the subtraction. Normalized HFA was significantly increased (relative to 0) during contralateral gripping in the set of ECoG sites showing highest gamma PLV in contralateral grip trials (CONTRA GRIP ECoG sites = blue line). Contralateral gripping resulted in significantly higher HFA than ipsilateral gripping in these sites (blue line in the left plot vs. blue line in the right plot). During contralateral gripping (left plot), HFA in CONTRA GRIP ECoG sites rose significantly faster compared to HFA in the other sites that showed highest coupling during ipsilateral gripping (IPSI GRIP ECoG sites = gray line). **B** The gamma phase difference between the two sets of cortical sites was offset negatively by approximately a quarter of a cycle during contralateral and positively during ipsilateral gripping. Only the latter was significant.

First, we found that contralateral grip trials resulted in significantly higher cortical HFA irrespective of bins (i.e. averaged across bins) compared to ipsilateral grip trials in the set of ECoG sites with highest gamma coupling during contralateral gripping (= CONTRA GRIP ECoG sites, P = 0.004, Wilcoxon signed-rank test). The difference in HFA between contra- and ipsilateral grip trials was less consistent in the ECoG sites with optimal coupling during ipsilateral gripping (P = 0.158, Wilcoxon signed-rank test). The blue line in the 2^nd^ row to the right in Figure 8A shows that during ipsilateral gripping, HFA activity didn’t even rise above baseline activity (obtained from the whole recording) in CONTRA GRIP ECoG sites (P = 0.535, t_27_=-0.6). Note that simultaneously, no significant STN spike-to-gamma coupling was present during ipsilateral gripping in these same sites (Figure 5A, bottom row).

Second, contralateral gripping resulted in different HFA profiles in the two distinct sets of cortical sites. HFA activity rose faster in the ECoG sites that showed highest coupling during contralateral gripping than in the other sites. (See Methods regarding other measures of HFA profile.)

The difference between the slopes in HFA increase was assessed by comparing the t-scores of the slope of a linear regression line fitted to each recording. Lines were fitted to the ascending period (−π:0 rad of the gamma cycle, p = 0.025, t_19_ = 2.6, only 20 recordings were included because the two ECoG sites were the same in the remaining eight recordings), as well as to the central and thus steepest 50% between the gamma trough and peak, located between −0.75π:−0.25π (p = 0.017, t_19_ = 2.4).

Both tests showed that the HFA increase was consistently steeper in the ECoG sites that showed the strongest coupling with STN spikes during contralateral gripping. This difference in HFA profile between ECoG sites was present only during contralateral but not during ipsilateral gripping.

Finally, we also tested whether the gamma phase was systematically shifted between the two sets of sites when either a contralateral or an ipsilateral grip was executed (within −0.1 – 0.4s window around movement onset). Figure 8B shows that the gamma phase was shifted by about a quarter of a cycle but with a change in sign between contra- and ipsilateral gripping: The average phase difference was −1.5 rad during contralateral gripping and 1.6 rad (95% CI = [0.7, 2.6]) during ipsilateral gripping. Only the latter was significant as the spread of phase differences was too large to estimate confidence intervals for the former.

## Discussion

We found that STN spike-to-cortical gamma phase coupling was strongest in motor cortical areas associated with hand movements, consistent with a link between subcortico-cortical gamma synchronisation and motor execution. Our comparison of trials with fast versus slow reaction times revealed that fast RTs were preceded by increased STN spike-to-cortical gamma phase coupling already at the time of the Go cue, suggesting a role for subcortico-cortical coupling in early motor programming. This relationship was only present for ECoG sites that showed the highest coupling during contralateral gripping but not in sites that were highly coupled with STN spikes during ipsilateral gripping. Thus, it is unlikely that increased STN spike-to-cortical gamma phase coupling alternatively reflected faster stimulus evaluation. A similar relationship between enhanced STN LFP to cortical gamma phase coupling and faster reaction times was recently reported in a dataset with overlapping subjects (Alhourani et al., 2018).

We also showed that STN spike probability was modulated relative to the gamma cycle during contralateral gripping. Considering the inhibitory effect that the STN exerts on the thalamus by exciting the inhibitory GPi/SNr, temporally focussed STN firing could enhance cortical gamma oscillations via rapidly oscillating downstream inhibition and disinhibition.

The gamma phase extracted at the time of STN spikes also was systematically offset between contra- and ipsilateral gripping, which provides further support for the hypothesis that the relative timing of STN spikes matters when executing distinct movements. During ipsilateral gripping, the timing of STN spikes was shifted by ~180° relative to the mean phase during contralateral gripping and thus occurred at the approximate opposite point of the gamma cycle. Presuming that the gamma cycle consists of alternating phases of relative depolarisation and relative hyperpolarisation of cortical neuronal populations, then this may be a mechanism to dynamically modify the functional consequences of STN spikes. Neurons in the GPi, a BG output nucleus, receive direct afferents from the STN and are also under cortical control via striatal direct-pathway medium spiny projection neurons. These striatal neurons inhibit the GPi. Hence, in the case of contralateral gripping, STN spikes may be timed such that they coincide exactly with these periods of inhibition. This would diminish the otherwise excitatory effect of STN spikes on GPi cells and result in thalamic and cortical disinhibition, correlates of which we saw as an increase in HFA during contralateral gripping (Figure 8A).

Alternatively, if the relative timing of STN spikes is offset by ~180°, they may arrive outside of periods of relative inhibition, which would help to maintain GPi firing and thus maintain tonic inhibition of the thalamus as should be the case during ipsilateral gripping.

Although many of the specifics of the present results are unique, several recent publications support the general idea that fine differences in the relative timing of neuronal activity in different components of the CBGTC loop may have a strong influence on the function of the circuit (Goldberg and Fee, 2012; Jantz et al., 2017). Precise temporal patterning of STN spikes thus could represent a key mechanism to avoid co-activation of cells that would otherwise activate competing effectors during movement execution. Loss of precision in temporal patterning might therefore lead to uncontrolled movements as seen in levodopa-induced dyskinesias of Parkinson’s disease or in Huntington’s chorea.

As expected, we found that cortical HFA, presumed to index local multi-unit activity, was higher during contralateral gripping. Additionally, we demonstrated that the HFA increased more steeply at those cortical sites where the gamma (60-80 Hz) phase was most strongly coupled with STN spikes during contralateral grip trials compared with the more dispersed set of sites, where coupling was strongest during ipsilateral gripping. If the phase of gamma oscillations is relevant for information separation between distinct cortical sites (Maris et al., 2013, 2016; van Ede et al., 2015), we would also expect a systematic phase offset between the gamma phase extracted from the two sets of sites. Indeed, the oscillations seemed to be shifted by one quarter of a cycle. This shift was only significant during ipsilateral gripping, where gamma power was relatively weak. Because a phase delay can also be interpreted as a phase advance of 360° minus the delay, the direction of the shift is ambiguous. Perhaps this offset facilitates separate processing of sensory information in the hemisphere ipsilateral to the gripping hand without increasing firing rates of “output units”. During contralateral gripping, where gamma power was high in both sets of sites, activity differed more in steepness of the HFA profile within the gamma cycle but not in a consistent gamma phase offset.

We also observed that the same cells that dynamically lock to gamma activity can lock to beta oscillations after movement completion. A rebound of post-movement beta oscillations has been associated with evaluation of a movement outcome. When feedback is as expected, the rebound is known to be high (Tan et al., 2014, 2016; Torrecillos et al., 2015). High spike-coupling to beta thus may consolidate the previous movement or bias against adaptation of the next motor response, although this remains to be tested. In this regard it is also interesting that if the initial cue indicated an ipsilateral grip, pre-movement cortical beta power was significantly elevated throughout the pre-Go signal delay period, consistent with an overall suppressive influence on contralateral output units prior to ipsilateral gripping. The variability of interspike intervals simultaneously decreased. Our findings support the emerging picture of two (although there may be more) distinct rhythms, the beta and the gamma rhythm, that can entrain neuronal spikes at different stages of motor control to facilitate or restrict motor output. In this light, gamma oscillations may reflect processing that is necessary for a change in motor state, such as the start of a movement or sudden cessation (Fischer et al., 2017).

Fluctuations in reaction times, but not in movement velocity or force, were linked to differences in gamma coupling. This seems at first surprising as local STN gamma power has been shown to correlate with movement velocity (Joundi et al., 2012; Lofredi et al., 2018). However, our task was not designed to capture a wide range of grip onset velocities and patients were studied during dopamine withdrawal, which reduces velocity-related gamma modulation (Lofredi et al., 2018).

We would like to highlight again that STN firing rates showed no consistent change across all recordings during the task. Thus, the timing of neuronal discharges with respect to cortical gamma phase was more predictive of movement than were cross-neuronal mean firing rates. Another indicator of a change in relative spike timing was the decrease of the interspike interval variability after the preparatory cue and in ipsilateral trials even during gripping.

Our study is limited in that ECoG coverage was not the same across patients and spanned limited areas of the cortical surface. However, we were able to perform group analyses by focussing on two sets of sites that showed the highest gamma coupling with contralateral and ipsilateral grips. We found spatially specific effects, such as the difference in coupling depending on reaction times and the phase-dependent spike probability modulation. These contrasts were unrelated to and did not depend on the ECoG contact pre-selection criterion and hence confirm that the selection procedure resulted in a physiologically relevant subset of ECoG sites involved in motor processing. The sites showing the highest gamma coupling during contralateral gripping were predominantly located precentral, in an area of motor cortex that seems to be involved in the control of upper limb movements (Cheyne et al., 2008). One site was located in lateral parietal area 5, which is also associated with direct corticospinal control of hand movements (Rathelot et al., 2017). However, we cannot exclude that coupling may also be present between STN spikes and other cortical sites that were not recorded.

Another caveat in this study is that firing rates in the STN were obtained from Parkinson’s disease patients. Firing rates in a subject at rest increase on average as the disease progresses (Remple et al., 2011). Whether our findings translate to neurologically healthy motor networks remains to be tested, although kinematics during task performance did not differ from those of subjects without a movement disorder (Kondylis et al. 2016). The coupling that we observed may be less specific than in healthy brains considering that progressive cell death in Parkinson’s disease has been associated with a loss of effector-specific activity (Bronfeld and Bar-Gad, 2011). Note that we included both single- and multi-unit recordings in the analysis. Considering that multiple cells need to be recruited in rhythmic firing to cause any downstream effects, inclusion of multi-unit recordings may even be beneficial when investigating spike-to-gamma phase coupling. Finally, our results are correlative and we cannot infer whether any electrophysiological correlates were actually causal for an observed behaviour, such as faster reaction times.

Taken together, our work provides intriguing evidence for the idea that the temporal structure of activity travelling through the basal ganglia is important for flexible motor control. Further investigation of information routing principles with multi-site recordings in the basal ganglia in simple movement tasks may provide essential new insights about basic mechanisms of motor control that can inform our ability to improve treatments for basal ganglia disorders.

## Acknowledgements

P. Fischer and P.B. are funded by the Medical Research Council (MC_UU_12024/1). R. M. R. was supported in part by the Walter L. Copeland Fund of The Pittsburgh Foundation. W. J. L. was supported by National Institute of Mental Health Grant R01MH107797 (PI, Avniel Ghuman; co-investigator, R. M. R.). P. Fries acknowledges grant support by DFG (SPP 1665, FOR 1847, FR2557/5-1-CORNET, FR2557/6-1-NeuroTMR), EU (HEALTH-F2-2008-200728-BrainSynch, FP7-604102-HBP, FP7-600730-Magnetrodes), a European Young Investigator Award, NIH (1U54MH091657-WU-Minn-Consortium-HCP), and LOEWE (NeFF).

## Author Contributions

W.L. and R.M.R. designed experiments and performed the recordings. W.J.N. reconstructed the ECoG positions in MNI space. P. Fischer analysed data and wrote the manuscript. W.J.N, R.S.T., P. Fries, P.B. and R.M.R. advised on analyses. All authors discussed the results and implications and commented on the manuscript at all stages.

## References

## Methods

### Subject details

The data recorded for this study were previously published elsewhere (Lipski et al., 2017b). 12 Parkinson’s disease patients performed a visually cued hand gripping task while undergoing deep brain stimulation surgery after overnight withdrawal from or a reduced dose of their dopaminergic medication. They provided written, informed consent in accordance with a protocol approved by the Institutional Review Board of the University of Pittsburgh (IRB Protocol no. PRO13110420). Demographic details are reported in Table S1.

### Method details

#### Task

The behavioural task was previously described (Kondylis et al., 2016; Lipski et al., 2017b). Patients had to perform unimanual left- or right-hand grip movements for at least 100ms with at least 10% of their previously determined maximal grip force. Considering that patients were currently undergoing deep brain stimulation surgery, they were not incentivised to respond as quickly as they could. Only trials with RTs longer than two seconds were treated as invalid. A single trial started with a yellow traffic light and an instructional cue either to the left or the right of it on a screen. The cue instructed participants whether to grip with the left or right hand. After a random interval ranging between 1-2s, the yellow light disappeared and a green or a red light came on. The green light instructed the patients to move and the red light served as NOGO cue and instructed patients to withhold the movement. The number of NOGO trials was variable across patients and not all patients performed a sufficient number of NOGO trials, thus NOGO trials were not analysed. The Go signal was presented on average 1.5 ±0.4s after the instructional cue. After trial completion, visual feedback was provided. The feedback was presented 1.9 ±0.4s after the Go signal and on average 0.43 ±0.41s after the grip was released. The average interval between the feedback and the next instructional cue was 1.4 ±0.3s (variable interval between 1-2s), so that on average every 4.7s ±1.2s a new trial started.

#### Electrophysiological recordings

STN microelectrode recordings were obtained with single glass-insulated platinum-iridium microelectrodes (FHC, Bowdoin ME) with impedances between 0.3 and 0.9 MΩ. Signals were filtered and amplified using the Guideline 4000 LP+ system (FHC; 125 Hz–20 kHz) and digitized at 30 kHz using the Grapevine Neural Interface System (NIPS; Ripple, Salt Lake City, UT). In one subject, the recordings were carried out using the Neuro-Omega recording system (Alpha Omega, Alpharetta, GA) using Parylene-insulated tungsten microelectrodes. Electrocorticography (ECoG) data was concurrently recorded with subdural strip electrodes placed temporarily via the existing burr hole used for DBS lead placement. The strips consisted of either 1 × 4, 1 × 6, or 1×8 (n = 11 patients; 2.3-mm exposed electrode diameter, 10-mm interelectrode distance) or a 2 × 14 [n = 1 patient; 1.2-mm exposed electrode diameter, 4-mm interelectrode distance; high-density (HD) contacts] platinum-iridium contacts (Ad-Tech Medical Instrument). ECoG signals were amplified, online notch filtered (at 60, 120, and 180 Hz), bandpass filtered (0.3–250 Hz) and digitized at 1,000 Hz using the Grapevine NIPS.

#### Data pre-processing

All analyses apart from the spike sorting were done in MATLAB (v. 2014b, The MathWorks Inc., Natick, Massachusetts, RRID:SCR_001622). Trials containing artefacts in the ECoG signal were removed after visual inspection, and only recordings with at least eight contralateral grip trials were included. This resulted in 28 recordings (12 single-unit recordings + 16 multi-unit recording) from 12 patients and an average number of 19 ± (SD) 8 contralateral and 18 ± (SD) 9 ipsilateral grip trials.

#### Spike sorting

The spike sorting procedure was described in detail elsewhere (Lipski et al., 2017a). Single- and multi-unit action potentials were identified offline using principal component analysis (Plexon, Dallas,TX). A cluster qualified as a single unit (SU) if: (1) the principal component cluster was clearly separated from other clusters associated with background activity and other units, (2) if it contained spike waveforms with a unimodal distribution in principal component space, and (3) if the inter-spike interval distribution showed a refractory period of at least 3ms (Starr et al., 2003; Schrock et al., 2009). For some SU recordings, the location of the principal component cluster drifted gradually, presumably due to a shift in distance between the electrode and the neuron caused by movement of the brain. Other recordings were classified as multi-unit (MU) recordings. In these, the principle components cluster seemed to include waveforms from multiple units, forming multimodal principal component distributions that could not be clearly separated, or that failed to meet the criterion of having a refractory period of 3ms.

### Quantification and statistical analysis

#### Time-frequency decomposition

Time-varying power and phase were obtained by band-pass filtering the data using Butterworth filters(4th order, two-pass, using fieldtrip functions_*ft_preproc_lowpassfilter* and *ft_preproc_highpassfilter* (Oostenveld et al., 2011)) and calculating the Hilbert transform. The time-frequency plots include the following frequency bins: 5-8 Hz for theta, 8-12 Hz for alpha, 12-20 Hz for low beta, 20-30 Hz for high beta followed by centre frequencies ranging from 40-150 Hz with a bin width of 20 Hz (incrementing by 10 Hz).

#### Event-related analyses of phase-coupling

Phase-locking values (PLV) can be obtained by calculating the length of the average of vectors, with each vector having a length of one on a unit circle and an angle corresponding to the ECoG phase (ϕ) coinciding with each STN spike (Lachaux et al., 2000). It was calculated as follows (n = number of spikes, ϕ_t_ = ECoG phase at the time of one spike):

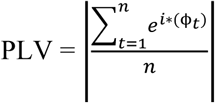

PLVs are bounded between 0 and 1, indicating zero or perfect phase coupling respectively. One important issue when calculating PLVs is that they are inflated when sample sizes are small. To correct for differences in sample size, the pairwise phase consistency (PPC) can be used instead as an unbiased estimator when a sufficiently large (>50) sample is available (Vinck et al., 2010; Aydore et al., 2013):

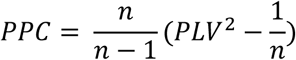

The PPC can attain negative values, because in the absence of phase locking, the expected PPC value for infinite amounts of data is zero, and the actual PPC values for limited amounts of data distribute around zero. Recordings with negative PPC values were excluded for analyses investigating phase offsets between contra- and ipsilateral grip trials as described below.

The fact that the firing rate of some cells increased or decreased around movement or cue onsets has important implications: If we would choose a fixed window size for calculating the PLV or PPC, we would base our analysis on variable and sometimes low numbers of spikes. Variable spike numbers would results in variable biases of the PLV metric, and low spike numbers in noisy estimates of the PPC metric. Another challenge was that some patients had substantially slower reaction times or longer grip durations than others because in the intraoperative setting, task constraints were not very strict.

To get comparable estimates of the coupling strength around each of the key trial events, we implemented the following custom-written event-locked variable-window width PLV estimation procedure: First, each interval between neighbouring task events was subdivided into 8 equidistant points (including the events) resulting in 29 time points for each trial (because the last point of an interval was identical to the first point of the next interval the points were concatenated the following way: [1-8, 2-8, 2-8, 2-8]). We had 4 intervals in total (Right/Left Cue to Go Cue, Go Cue to Grip Onset, Grip Onset to Grip Offset, Grip Offset until the end of a 0.4s long post-movement window). The first time point of each trial was set at the onset of the Right/Left Cue and the final one at 0.4 seconds after the grip offset. This procedure ensures that a given plot shows unbiased estimates of the coupling strength around all key task events and not just for one single event, which would be the case if it was locked only to movement or cue onsets.

To include the same number of spikes around each time point, and thus avoid fluctuations of the PLV just because of differences in sample size, the windows centred around each time point were scaled in width. Separately for each recording, the target spike number was set to the average number of spikes occurring within a 0.3s window (obtained by summing up all spikes across all trials, dividing by the total trial length and multiplying by 0.3s). If this number was below 50, it was set to 50 to avoid very small and thus less representative samples. Each window was placed symmetrically around each time point and its width was set such that the sum of spikes across all trials would match the target number as closely as possible (Figure S8). This procedure allowed an unbiased comparison of coupling over time despite changes in firing rate that occurred in some recordings. The average window size was 0.32 ± (SD) 0.04s (min.: 0.16s, max.: 0.75s). Note that the procedure did not ensure that the number of spikes was the same across subjects. Ensuring this would have required an even larger variability of window widths. The average number of spikes per window was 171 ± (SD) 121 spikes. However, our main results hold both for PLV and PPC values, with the latter eliminating biases resulting from different sample sizes.

Each matrix was composed of 16 frequency bins and 29 time points. The resulting 28 matrices (one for each of the 28 recordings) were event-locked to all four task events and could thus be easily averaged across recordings despite differences in reaction times or grip duration. The coarsely sampled matrices were smoothed using the MATLAB function *interp2* with an interpolation factor of 2 resulting in a grid with a resolution of 61 * 133. Finally, the time-dimension was resized using the MATLAB function *imresize* to rescale the inter-event intervals to the average recorded durations.

#### Firing characteristics

Firing rates were calculated within a 300ms window around each time point to get a time-evolving estimate. Three additional firing characteristics were calculated similar as in Rule et al. (2017): The interspike interval coefficient of variation (ISI CV), the percentage of bursting (%burst) and the mode of the ISI (ISI mode). Because properties such as the coefficient of variation (CV) can again be biased by small sample sizes and to enable a direct comparison, we used the same window widths to calculate time-evolving firing characteristics as outlined above for calculating the time-evolving PLV. A robust version of the ISI CV was calculated as the median absolute deviation of all ISIs divided by the median and multiplied by 100. The %burst metric was defined as the percentage of all ISIs shorter than 10ms. To calculate the ISI mode, we first excluded all bursts, and then computed a probability density estimate (using the MATLAB function *ksdensity*) of which the mode was extracted. The baseline firing characteristics that were used to test for significant deviations were computed by splitting the whole recording into segments with the same number of spikes as were included when computing it across trials or into 300ms long segments for computing the baseline firing rates. To get a robust estimate of individual baseline values, the median of all segments was computed for each individual recording.

#### ECoG contact position normalization to MNI space

The temporarily placed subdural ECoG strips were localized by aligning intraoperative fluoroscopy (x-ray) images of the strip to co-registered preoperative MRI and postoperative CT scans (Randazzo et al., 2016). In brief, preoperative MRI scans were co-registered to postoperative CT using SPM (http://www.fil.ion.ucl.ac.uk/spm/software/spm12/) and consecutively decomposed as 3D skull and brain surfaces using Freesurfer (Dale et al., 1999; https://surfer.nmr.mgh.harvard.edu/). The reconstructions were then co-registered to common landmarks visible in the intraoperative fluoroscopy images (stereotactic frame pins, implanted DBS electrodes and skull outline). The parallax effect of the fluoroscopic images was accounted for by using the measured distance from the radiation source to the subject’s skull to adjust the projection of the skull/brain surfaces. Three-dimensional coordinates of ECoG recording sites were obtained in native Freesurfer space, based on the alignment of the ECoG strip in the fluoroscopic image to the cortical surface reconstruction. To allow group analysis, the electrode locations were then transformed to ICBM152 population averaged cortex space using population-based normalization of the native surface reconstructions (Saad and Reynolds, 2012).

#### ECoG contact selection

To get a local estimate of the cortical power and phase, we computed bipolar configurations by subtracting neighbouring ECoG contacts resulting in n-1 bipolar channels if n contacts were recorded. The ECoG contact with the highest PLV between 60-80 Hz and within a −0.1 – 0.4s around movement onset during contralateral grip trials were pre-selected for further analyses. 60-80 Hz was chosen, as previous research has shown coherence between STN LFP and cortical oscillations in this frequency band (Williams et al., 2002; Litvak et al., 2012). We also performed the same selection procedure to find the contacts showing the highest coupling during ipsilateral grip trials.

#### Topography of gamma coupling and power

In the topoplots of movement-related gamma and beta power changes, the average across all contacts was displayed. Relative power from a −0.1 – 0.4s window around grip onsets was normalized by the average power from the same contact across the whole recording. All points of the cortical mesh within a 4 mm radius (to increase spatial specificity when including all contacts) from the corresponding ECoG site were coloured uniformly with the corresponding value and then averaged.

To illustrate the cortical distribution of gamma coupling during contra- and ipsilateral gripping, we displayed the two sets of ECoG sites with highest gamma coupling during contralateral gripping as well as during ipsilateral gripping. We also show the distribution of the subset of contacts displaying significant movement-related spike-to-60-80 Hz gamma coupling in the same window. Significance was assessed again by creating permutation distributions. To pass the test, two conditions had to be met: First, the original PPC should be significantly higher than the null-distribution created by randomly permuting the ECoG LFP-to-STN spike association 500 times across trials. With this method, the pattern of spikes and the more pronounced gamma synchrony in the ECoG LFP at movement onset remains intact. Second, the original PPC should be significantly higher around movement onset compared to a pre-movement period (spikes were drawn from a window ranging from −3 to −2s before movement onset and matched in the number of spikes) from where spikes and coincident ECoG phases were extracted. We applied one-sided tests (α = 0.1, but both conditions had to be significant) such that a larger number of contacts, which is more representative of the overall distribution, was displayed in the topoplot. Cortical mesh areas within a 7 mm radius around each bipolar contact site (defined as the average location between the two contacts used for the bipolar calculation) were displayed as significant to ensure good visibility of each single contact.

#### Cluster-based permutation procedure

Significance tests in plots showing multiple time- or frequency points were performed using a cluster-based permutation procedure for multiple-comparison correction (Maris and Oostenveld, 2007). Condition labels of the original samples (e.g. median-split fast or slow RTs) were randomly permuted 1000 times such that for each recording the order of subtraction can change. When firing rates in individual recordings were compared relative to a baseline period (ranging from −0.5s before the Left/Right cue to the Go signal), the normalized firing rate was compared against 0 by flipping the sign of the firing rates of a subset of randomly chosen trials. This permutation procedure results in a null-hypothesis distribution of 1000 differences. Suprathreshold-clusters (pre-cluster threshold: P<0.05) were obtained for the original unpermuted data and for each permutation sample by computing the z-scores relative to the permutation distribution. If the absolute sum of the z-scores within the original suprathreshold-clusters exceeded the 95^th^ percentile of the 1000 largest absolute sums of z-scores from the permutation distribution, it was considered statistically significant.

To test if PLVs were significantly elevated, we created a permutation distribution by shuffling the LFP-to-spike relationship in each recording 500 times. The same windows were chosen as for the calculation of the original PLVs but the trial-association between the ECoG LFP and STN spikes was permuted such that, for example, the spike train from trial 1 was paired with the LFP from trial 4, resulting in phases that could have been observed by chance if no spike-to-phase coupling existed. Importantly, the same shuffling was applied to all time- and frequency points within each of the 500 permutations. If different randomizations were applied for each time- or frequency point, the natural appearance of clusters would be prevented, and the test, which is again based on the z-scores of the largest suprathreshold clusters as above, would not be sufficiently conservative.

Whenever time-frequency plots were compared between contra- and ipsilateral grip trials, the cluster-based permutation procedure was performed on the differences (e.g. between PLVs during fast vs. slow RTs or between original PLVs and shuffled PLVs).

#### Phase-dependent spike probability and phase flipping according to local high-frequency activity

Average PLVs can be high across recordings although the average preferred phase could differ from recording to recording. When computing the PLV, the information of preferred phase is not retained as only the vector length of the average phase vector is taken into account, reflecting how strongly bundled or spread the phase values are. We intended to test, whether spikes were consistently more probable at certain phases and less probable at others.

To examine if the spike probability was consistently increased or decreased at certain phases, we subdivided the cortical gamma signal into five phase bins to calculate the percentage of spikes occurring within these bins. An issue that needs to be dealt with beforehand is that the phase of the cortical oscillation can be polarity-reversed depending on the source of the gamma oscillation and order of subtraction of the two channels for the bipolar configuration. Gamma oscillations are thought to reflect fluctuations in local excitability. To ensure that the gamma phase has the same polarity across all recordings, we performed the following automatized phase-flipping procedure:

First, the local high-frequency activity (HFA) was computed by high-pass filtering the ECoG signal at 300 Hz, full-wave rectifying and low-pass filtering at 100 Hz. To increase the signal-to-noise ratio, the whole recording was used. Next, we extracted the 60-80 Hz gamma phase from the same signal and subdivided it into 126 equally spaced, non-overlapping phase bins with a bin width of 0.05π. For each bin, the average HFA activity was computed. The resulting 126 points-long vector was smoothed with a running average (width = 20 samples). To check if the gamma phase should be flipped, the average HFA was calculated within a 0.4π wide window centred at the middle (at 0π of the gamma phase, which is where the gamma-filtered data would have its peak) and an average of the two pieces on the side (0.2π wide on each side, where the gamma-filtered data would have its minima). If the average HFA from the middle was lower than the average HFA from the sides, the gamma phase was flipped (see Figure S7 A). All cortical recordings showed a distinguishable peak, and thus this method helped to ensure that the cortical gamma phase was flipped such that the gamma peak consistently matched the HFA.

To compare changes in spike probability across the gamma cycle, we split the gamma phase into five non-overlapping phase-bins resulting in bins with a width of 0.4π. Five bins were chosen to allow for a resolution that does not result in too many multiple comparisons but is still high enough to detect differences.

We computed movement-related spike probabilities for the five phase bins based on data from a - 0.1 – 0.4s window around movement onset and baseline spike probabilities from a same sized window starting 3s before movement onset. The relative changes were obtained by subtracting baseline spike probabilities from movement-related spike probabilities.

Two-factorial 5 (bin) x 2 (effector side) repeated-measures ANOVAs were computed in SPSS (IBM Statistics SPSS 22, RRID:SCR_002865), to compare spike probabilities across the five phase-bins. Q-Q plots of the residuals were visually inspected to exclude strong deviations from normality. If the sphericity assumption was violated, Huynh-Feldt correction was applied. Subsequent pairwise comparisons were performed using t-tests or Wilcoxon signed-rank tests if the normality assumption (assessed by Lilliefors tests) was violated.

#### Comparison of phase offsets between contra- and ipsilateral gripping

Another key question was whether the timing of STN spikes relative to the cortical gamma phase systematically differed between contra- and ipsilateral grip trials. We computed the circular median of all phases coinciding with STN spikes around movement onset (in a −0.1 – 0.4s window). Only recordings with positive PPC values were included resulting in n=23 as an estimate of the average phase would not be meaningful if the phases were relatively uniformly distributed. The angle difference between contra- and ipsilateral gripping was first tested against zero by computing confidence intervals based on the 23 angle differences (*circ_confmean* function from the *circ_stats* toolbox (Berens, 2009)). We decided not to scale the weight of the angles by the number of spikes included or the PPC strength for each recording as this would bias the average across all recordings towards few data points. Spike numbers were not subsampled, which could have been a step to match spike numbers between recordings, as the estimate of the mean angle only gets more precise with a larger number of representative samples. We also investigated whether the offset of 180° degrees, which became apparent, was specific in time by plotting the time course of the p-values of a V-test (*circ_vtest* function from the *circ_stats* toolbox (Berens, 2009)). A V-test examines the phase differences for circular uniformity, similarly as a Rayleigh test, but is more powerful because a mean difference – in this case 180° – can be specified.

#### Temporal profile of cortical high-frequency activity evolving with the gamma cycle

As we saw that the modulation of STN spike timing relative to the cortical gamma phase was distinct for different sets of cortical sites and presumed that this somewhat interacts with local cortical processing, a next logical step was to ask if the profile of the cortical high-frequency activity that fluctuates with each gamma cycle was also distinct. For 8 of the 28 recordings, the ECoG sites with highest gamma coupling for contra- and ipsilateral grip trials were identical, and thus were excluded from this analysis resulting in n=20. Cortical HFA (see “Phase-dependent spike probability and phase flipping according to local high-frequency activity” section above) extracted from −0.1 – 0.4s around movement onset and normalized by the mean HFA across the whole recording was plotted separately for contra- and ipsilateral grip trials and the two sets of ECoG sites. As the relative increase in HFA activity differed between the two sets of sites, which may merely be related to the distance of the source from the recording site or impedance of the tissue, we subtracted the average (across both grip trials) of each site from each of the displayed lines. We also tested whether the slope of the HFA increase differed between the two sets of sites.

We compared the slopes of the increase instead of the peak values because HFA levels appeared to be biased by the proximity of the recording sites to the cells generating the HFA or by differences in tissue impedance even when normalized by the session mean (see Fig. 7a, second row, right). Different levels of baseline activity, i.e. a general offset, would not affect the slope of the increase. Calculating the relative modulation depth would be another alternative. However, this option is inferior to comparing the slopes because the precise location of the trough and peak may vary, which would introduce more variability.

To calculate the slopes, the HFA was iteratively smoothed first with a running average of 40 (of128) samples, and then with a running average of 20 samples resulting in smooth activity traces. Then, the t-scores of the slope of a linear regression line fitted to the ascending phase of the gamma cycle (between −π:0 rad, and between −0.75π:−0.25π rad to assess only the steepest part) were computed for each recording to evaluate them on the group level. The difference in gamma phase between the two sets of ECoG sites during contralateral and ipsilateral gripping (−0.1 – 0.4s around movement onset) was also analysed.

### Data and software availability

The datasets and code generated during the current study are available on reasonable request.

